# Biosynthesis of glucosaminyl phosphatidylglycerol in *Pseudomonas aeruginosa*

**DOI:** 10.1101/2024.10.10.617631

**Authors:** Fabiha Zaheen Khan, Kelli L. Palmer, Ziqiang Guan

## Abstract

Glucosaminyl phosphatidylglycerol (GlcN-PG) was first identified in bacteria in the 1960s and was recently reported in *Pseudomonas aeruginosa*. Despite the important implications in altering membrane charge (by the modification of anionic PG with cationic glucosamine), the biosynthesis and functions of GlcN-PG have remained uncharacterized. Using bioinformatic and lipidomic analysis, we identified a 3-gene predicted operon, renamed as *gpgSDF*, that is responsible for the biosynthesis and potential transport of GlcN-PG in *P. aeruginosa*: *gpgS* encodes a novel glycotransferase that is responsible for the modification of phosphatidylglycerol (PG) with *N*-acetylglucosamine (GlcNAc) to produce GlcNAc-PG, and *gpgD* encodes a novel deacetylase that removes the acetyl group from GlcNAc-PG to produce GlcN-PG. The third gene, *gpgF*, is predicted to encode a flippase whose activity remains to be experimentally verified. A *P. aeruginosa gpgD* transposon mutant accumulates GlcNAc-PG and lacks GlcN-PG, and as expected, the complementation of *gpgD* restores the production of GlcN-PG. Moreover, the heterologous expression of *gpgSDF* in *Escherichia coli* resulted in production of both GlcNAc-PG and GlcN-PG. The identification of the biosynthetic genes of GlcN-PG paves the way for the investigation of its biological and pathological functions, which has significant implications in our understanding of the unique membrane physiology, pathogenesis and antimicrobial resistance of *P. aeruginosa*.

**Importance:** The identification of the biosynthetic genes of GlcN-PG paves the way for the investigation of its biological and pathological functions, which has significant implications in our understanding of the unique membrane physiology, pathogenesis and antimicrobial resistance of *P. aeruginosa*.

## Introduction

*Pseudomonas aeruginosa* is a major Gram-negative pathogen widely responsible for pneumonia, surgical infection, bacteremia, and other life-threatening infections in immunocompromised individuals with underlying diseases such as cystic fibrosis (CF) (1, 2) and cancer (3). *P. aeruginosa* possesses an unusual ability to colonize diverse environments and rapidly develop antibiotic resistance. Multi-drug-resistant (MDR) and extreme drug-resistant (XDR) high-risk strains are widespread in healthcare settings, making the treatment of certain *P. aeruginosa* infections extremely challenging (4–6).

An important mechanism for bacteria to cope with antimicrobial stresses and develop antibiotic resistance is by altering the charge of their membrane lipids. A highly conserved and perhaps most studied enzyme responsible for modifying lipids in bacteria is MprF (multiple peptide resistance factor), which modifies lipids via the transfer of amino acids from charged tRNAs to the head groups of anionic phosphatidylglycerol (PG) and cardiolipin (CL) (7, 8), as well as to neutral glycolipids (9, 10). *P. aeruginosa* encodes MprF and produces alanine-modified PG (11). Additionally in *P. aeruginosa*, modifications of the phosphate groups of lipid A (the lipid anchor of lipopolysaccharide) with positively charged moieties such as aminoarabinose and phosphoethanolamine are critical for conferring resistance to cationic antimicrobial peptides and polymyxin (2, 12–16).

The modification of PG with cationic glucosamine, producing zwitterionic glucosaminyl phosphatidylglycerol (GlcN-PG), was recently reported in *P. aeruginosa* (17). Despite the obvious implications in altering membrane charge, and the fact that GlcN-PG was first observed in various bacteria in the 1960s (18–21), the biosynthesis and functions of GlcN-PG have remained uncharacterized.

In this study, by combining bioinformatic, genetic, and lipidomic approaches, we identified the biosynthetic genes required for GlcN-PG synthesis in *P. aeruginosa*. Using the PA14 strain as a model, we show that a 3-gene predicted operon (consisting of PA14_56030, PA14_56040 and PA14_56050) is required for the synthesis and possible transport of GlcN-PG. We have thus renamed this operon as *gpgSDF* (<u>G</u>lcN-<u>PG</u> <u>s</u>ynthase, <u>d</u>eacetylase and flippase). Since *gpgSDF* is conserved in other strains of *P. aeruginosa*, this work has significant implications for understanding the mechanisms of physiology and pathogenesis of *P. aeruginosa*.

## Experimental Procedures

### Bacterial Strains and Routine Growth Conditions

The bacterial strains used in the study are described in Table 1. For lipidomic analysis, bacterial cells were grown in Luria Bertani (LB) broth (10g/L tryptone, 5g/L yeast extract, 5g/L NaCl) at 37 °C with shaking at 220-250 rpm for 18-20 hours. The optical density (OD) was measured in a disposable cuvette (Thermo Fisher Scientific, Waltham, MA) as absorption at 600 nm by a spectrophotometer (Thermo Scientific Genesys 30).

**Table 1.** Bacterial Stains and Plasmid Used in This Study.

| Species and strains | Description | Source or reference |
| --- | --- | --- |
| *P. aeruginosa* |  |  |
| PAO1 | Wild type (first isolated in 1954 from a wound in Melbourne, Australia) | (54, 55) |
| PA14 | Wild type (first isolated in early 1970s from the blood of a burn patient at Mercy Hospital in Pittsburg, PA, USA) | (56) |
| PA103 (ATCC 29260) | Wild type | (57) |
| ATCC 27853_NP446 | Wild type (isolated in 1971 from a blood specimen in Peter Bent Brigham Hospital, Boston, USA) | (58) |
| PA14_56030::Mar2xT7<br>(*gpgS*::Tn) | PA14 with Mar2xT7 Tn insertion at nucleotide (nt) position 896 of 1116 nt PA14_56030 (*gpgS*) locus | (22) |
| PA14_56040::Mar2xT7<br>(*gpgD*::Tn) | PA14 with Mar2xT7 Tn insertion at nt position 226 of 762 nt PA14_56040 (*gpgD*) locus | (22) |
| PA14_56050::MarxT7<br>(*gpgF*::Tn) | PA14 with Mar2xT7 Tn insertion at nt position 403 of 1002 nt PA14_56050 (*gpgF*) locus | (22) |
| PA14_56040::Mar2xT7(pMQ72-Tet-*gpgD*) | PA14_56040::Mar2xT7 with the arabinose-inducible complementation vector pMQ72-Tet-*gpgD* | This work |
| *E. coli* |  |  |
| *E. coli* DH5α | Cloning host strain | Thermo Fisher Scientific |
| DH5α (pWO-GEM-T) | pWO-GEM-T with *gpgSDF* predicted operon | This work |

The *P. aeruginosa* PA14 MAR2xT7 transposon mutants carrying a gentamicin resistance cassette were previously reported (22) and were grown in LB broth containing 15 μg/ml gentamicin. *Escherichia coli* DH5α (Thermo Fisher Scientific) with the plasmid pGEM-T (Promega PR-A3600) was grown in LB broth containing 50 μg/ml ampicillin.

### Complementation of the *gpgD* transposon mutant of PA14

The PA14 *gpgD* gene was amplified using primers 1 and 2 (Table S1), which introduced regions complementary to the plasmid pMQ72 (23) (obtained from Dr. Peter Jorth). The pMQ72 plasmid was digested using EcoRI and HindIII. The *gpgD* amplicon and digested pMQ72 were used for Gibson assembly. The reaction was transformed into chemically competent *E. coli* DH5α cells. Gentamicin at 30 µg/mL was used for selection of pMQ72 in *E. coli*. The pMQ72-*gpgD* plasmid was extracted using the Thermo Scientific GeneJET Plasmid Extraction Kit following the manufacturer’s protocol, and the *gpgD* sequence was confirmed by Sanger sequencing at the University of Texas at Dallas Genome Center using primer pairs 3 and 4, 5 and 6, 7 and 8, and 9 and 10 (Table S1). After confirmation, the pMQ72-*gpgD* plasmid was amplified using primers 11 and 12 (Table S1) to generate a linear fragment that excluded the gentamicin resistance cassette. The tetracycline resistance cassette from pEX18Tc (24) was amplified using primers 13 and 14, which introduced overhangs corresponding to pMQ72 (Table S1). The pMQ72 fragment and the tetracycline resistance cassette were assembled as described above. Successful insertion of the tetracycline cassette was confirmed using primers 15 and 16 (Table S1). The resulting plasmid, pMQ72-Tet-*gpgD*, was chemically transformed into *E. coli* S17 cells (obtained from Dr. Trusha Parekh), and transformants were selected with tetracycline at 10 µg/mL. The S17 transformants were replica-plated on LB plates with 30 µg/mL gentamicin, where no growth was observed, as expected. The pMQ72-Tet-*gpgD* plasmid was conjugated into PA14 *gpgD*::Tn cells using *E. coli* S17 as the donor. Transconjugant colonies were selected on Vogel-Bonner minimal medium (VBMM) agar with 100 µg/mL ampicillin and 100 µg/mL tetracycline (25). Transconjugant colonies were confirmed by colony PCR using primer pairs 3 and 10, and 15 and 16 (Table S1). To induce expression of *gpgD* from the *araBAD* promoter of pMQ72, PA14 *gpgD*::Tn(pMQ72-Tet-*gpgD*) were grown to late exponential phase in LB (OD_600_∼1.0) with 100 µg/mL tetracycline +/- 1% L-arabinose (26). Cells were pelleted for acidic Bligh-Dyer lipid extractions and normal-phase LC/MS analysis.

### Heterologous expression of *gpgSDF* in *E. coli*

The *gpgSDF* predicted operon was amplified by PCR using specific primers (Table S1) and cloned into pGEM-T (27) by TA cloning per the manufacturer’s instructions. The newly assembled pWO-GEM-T plasmid was confirmed through PCR reactions with T7 and SP6 and sequencing primer pairs with agarose gel electrophoresis analysis of products. The plasmid was transformed into *E. coli* DH5α and maintained by growth in the presence of 50 μg/ml of ampicillin. Cells were pelleted for acidic Bligh-Dyer lipid extractions and normal-phase LC/MS analysis.

### Acidic Bligh-Dyer Lipid Extraction

The acidic Bligh-Dyer lipid extractions were performed as previously described (9, 28). Briefly, bacterial cells were grown in 20 mL of LB broth for ∼20 hours; 19 mL of each culture was pelleted at 7000 rpm for 10 min. The cell pellet was washed twice with 1X phosphate buffer saline (PBS, Sigma-Aldrich, St. Louis, MO). Each cell pellet was resuspended in 0.8 mL of 1X PBS and transferred to a 17-mL glass tube with a Teflon-lined cap (Corning Pyrex^TM^, VWR, Radnor, PA). Afterwards, 1 mL of chloroform and 2 mL of methanol were added to generate a single-phase Bligh-Dyer solution. This solution was incubated for 20 min at room temperature with intermittent mixing. After centrifugation at 1700 rpm for 10 min, the supernatant was transferred to a new glass tube, which was followed by the addition of 100 μl HCl (37%), 0.9 mL 1X PBS and 1 mL chloroform to generate a two-phase Bligh-Dyer solution. After mixing by a vortex, the solution was separated into two phases by centrifugation at 1700 rpm for 5 min at room temperature. The lower phase was recovered and dried under a stream of nitrogen gas before being stored at −80°C prior to lipidomic analysis.

### Normal Phase LC/MS/MS Analysis

Lipidomic analysis by normal phase LC/MS/MS was described previously (9, 28). Briefly, normal phase LC was performed on an Agilent 1200 Quaternary LC system equipped with an Ascentis Silica HPLC column, 5 μm, 25 cm x 2.1 mm (Sigma-Aldrich, St. Louis, MO). Mobile phase A consisted of chloroform/methanol/aqueous ammonium hydroxide (800:195:5, v/v); mobile phase B consisted of chloroform/methanol/water/aqueous ammonium hydroxide (600:340:50:5, v/v); mobile phase C consisted of chloroform/methanol/water/aqueous ammonium hydroxide (450:450:95:5, v/v). The elution program consisted of the following: 100% mobile phase A was held isocratically for 2 min and then linearly increased to 100% mobile phase B over 14 min and held at 100% B for 11 min. The LC gradient was then changed to 100% mobile phase C over 3 min and held at 100% C for 3 min, and finally returned to 100% A over 0.5 min and held at 100% A for 5 min. The LC eluent (with a total flow rate of 300 μl/min) was introduced into the ESI source of a high resolution TripleTOF5600 mass spectrometer (Sciex, Framingham, MA). The instrumental settings for negative ion ESI/MS and MS/MS analysis of lipid species were as follows: IS = -4500 V; CUR = 20 psi; GSI = 20 psi; DP = -55 V; and FP = -150 V. The MS/MS analysis used nitrogen as the collision gas. Analyst TF1.5 software (Sciex, Framingham, MA) was used for data analysis. All analysis was performed by at least duplicate experiments.

## Results

### Identification of GlcN-PG in *P. aeruginosa* by LC/MS/MS

The lipid extract of *P. aeruginosa* strain PA14 cells was analyzed by normal phase LC/MS in both negative ion and positive ion modes. As shown by the negative total ion chromatogram (TIC) (Fig. 1A), the major detected lipids include alkyl quinolones (AQs), diacylglycerol (DAG), phosphatidylglycerol (PG), phosphatidylethanolamine (PE), cardiolipin (CL) and phosphatidylcholine (PC). These common lipids and their biosynthesis have been extensively studied in *P. aeruginosa* and other bacteria (29–31).

**Figure 1.**
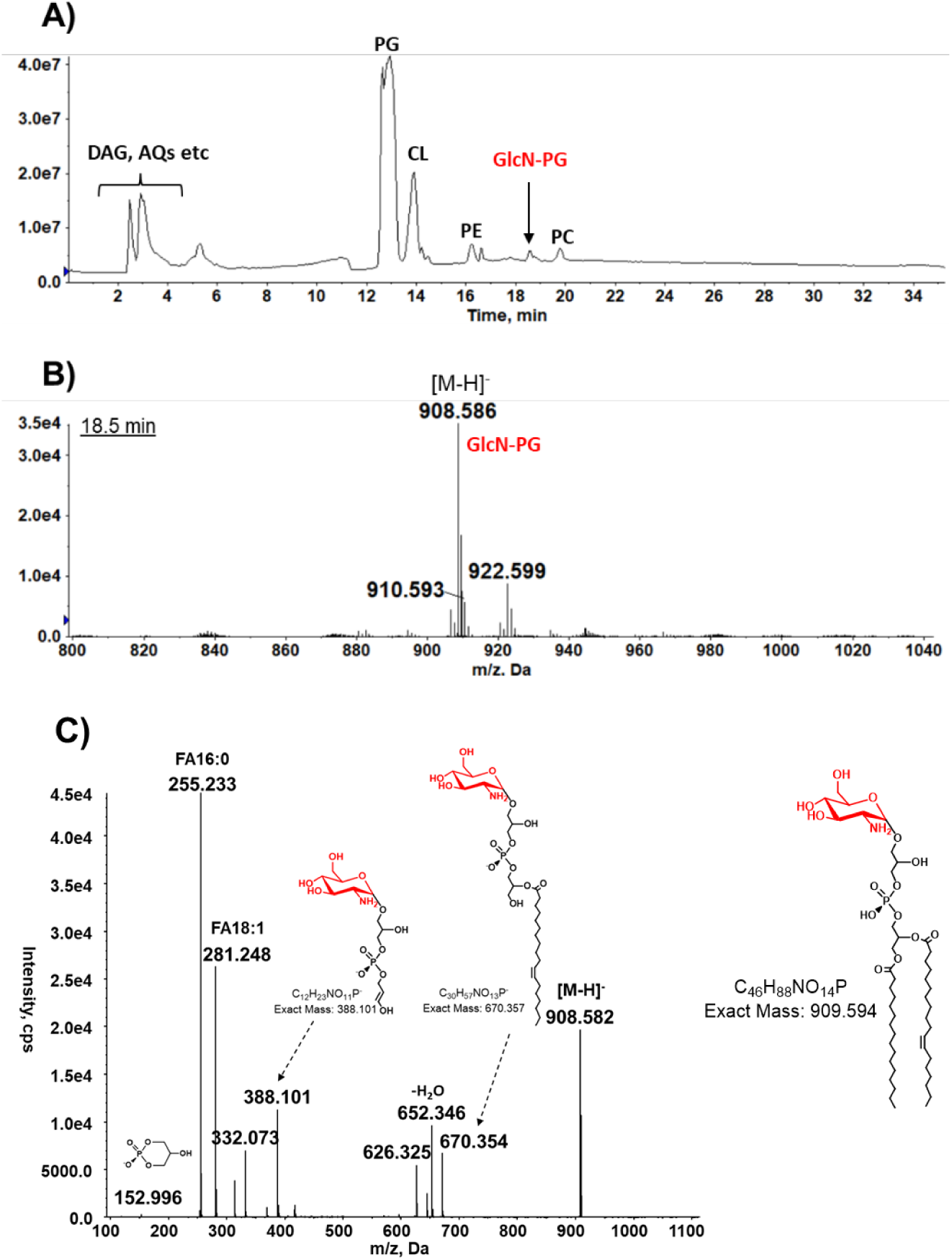
Identification of GlcN-PG in *P. aeruginosa* PA14 by LC/MS/MS. The lipid extract of PA14 strain is analyzed by normal phase LC/MS/MS, with GlcN-PG being identified by exact mass measurement and MS/MS. A) Negative total ion chromatogram (TIC) of normal phase LC/MS of the lipid extract of *P. aeruginosa* strain PA14. B) Negative ion ESI mass spectrum (at ∼18.5 min) showing the [M-H]^-^ ions of GlcN-PG. The observed exact mass (908.586) matches with the predicted exact mass (908.587) for the [M-H]^-^ ion of GlcN-PG (16:0/18:1). C) MS/MS of [M-H]^-^ ion at *m/z* 908.5 produces fragment ions consistent with GlcN-PG (16:0/18:1) whose chemical structure is shown. The linkage positions of fatty acids and GlcN as depicted in the chemical structure of GlcN-PG (Fig. 1C) have not been determined experimentally and are for illustrative purposes.

An uncommon lipid with a major [M-H]^-^ molecular ion at *m/z* 908.5 was eluted at ∼18.5 min (Fig. 1B). Based on tandem mass spectrometry (MS/MS) (Fig. 1C) and exact mass measurement, it was identified as glucosaminyl-phosphatidylglycerol (GlcN-PG) recently reported in *P. aeruginosa* (17). GlcN-PG was previously found in other bacteria (18, 19, 21, 32). To our knowledge, there have been no reports on the studies of the biosynthetic genes and functions of GlcN-PG in *P. aeruginosa* or any other bacteria.

### Identification of the biosynthetic genes of GlcN-PG

Enzymatically, GlcN-PG is most likely produced from PG via the covalent modification with GlcN. Indeed, this was supported by the observation that during the pellicle growth of *P. aeruginosa*, the increase of GlcN-PG was accompanied by the decrease of PG (17). Previously it was also observed in *Bacillus megaterium* that GlcN-PG was increased, while PG was decreased, when cells were grown in acidic conditions (32).

Given the biochemical mechanisms of other amino sugar modifications (33, 34), we hypothesize that GlcN-PG is likely produced from its immediate precursor, GlcNAc-PG, by *N*-deacetylation, an enzymatic process that is known to be involved in several biosynthetic pathways. For examples, LpxC is the UDP-3-*O*-(*R*-3-hydroxymyristoyl)-*N*-acetylglucosamine deacetylase required for the biosynthesis of lipid A, the lipid anchor of lipopolysaccharide (LPS) in Gram-negative bacteria (35–37). Deacetylation mediated by carbohydrate esterase family 4 proteins (38) plays a vital role in exopolysaccharide processing in different bacterial species, including *P. aeruginosa*.

The potential involvement of deacetylation in the biosynthetic process of GlcN-PG prompted us to carefully examine the PA14 lipidomic data and identify a low level of GlcNAc-PG (Fig. 2A). The [M-H]^-^ ion of GlcNAc-PG is observed at *m/z* 950.59, corresponding to the addition of an acetyl group (42 Da) to GlcN-PG (*m/z* 908.58). The detection of GlcNAc-PG provides an important clue for the existence of a deacetylase that converts GlcNAc-PG to GlcN-PG.

**Figure 2.**
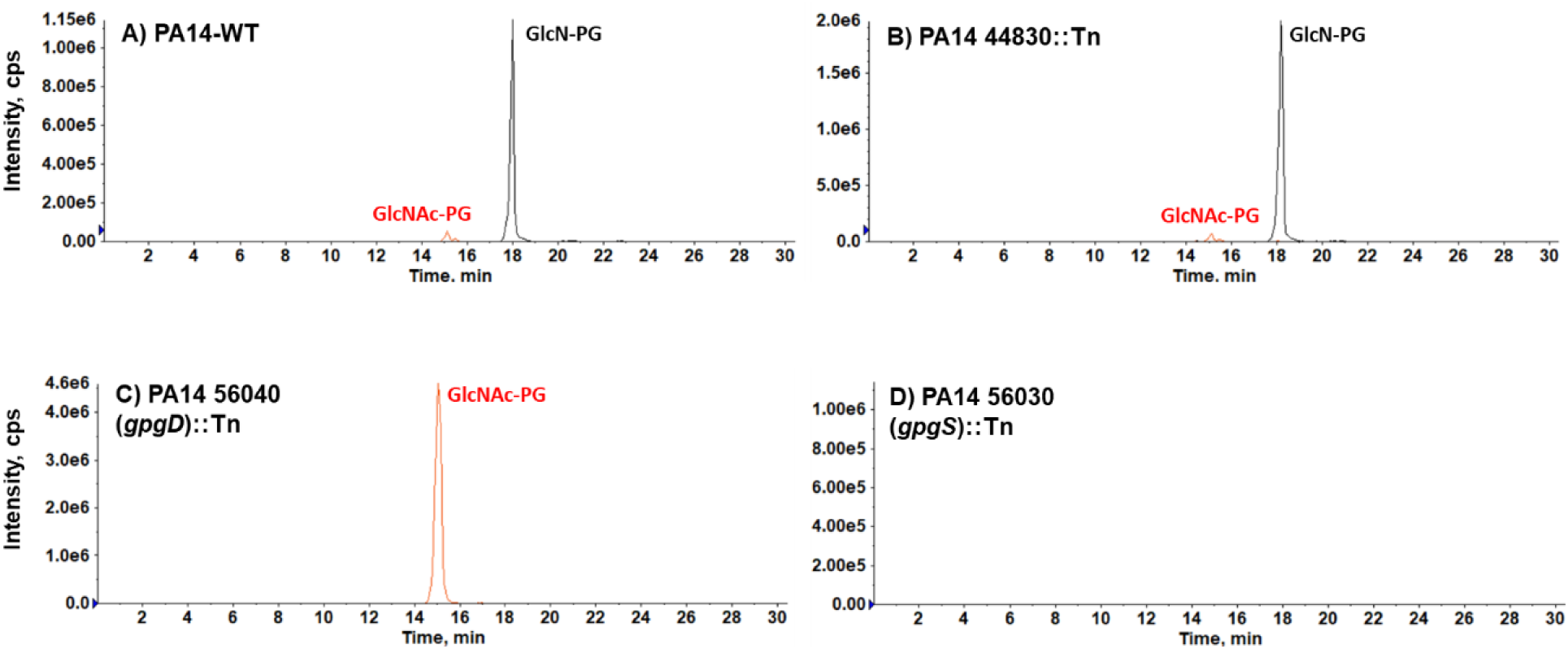
PA14_56030 (*gpgS*) and PA14_56040 (*gpgD*) genes are required for the synthesis of GlcNAc-PG and GlcN-PG in *P. aeruginosa*. LC/MS analysis of the transposon mutants of PA14_56030 (*gpgS*, putative glycotransferase) and PA14_56040 (*gpgD*, putative deacetylase) confirm that they are responsible for the synthesis of GlcNAc-PG and GlcN-PG in *P. aeruginosa,* respectively. A, B) GlcN-PG and a low level of GlcNAc-PG are present in the wild-type PA14 and PA14_44830::Tn mutant. C) GlcN-PG is depleted, and GlcNAc-PG is drastically accumulated in the PA14_56040::Tn mutant. D) Both GlcN-PG and GlcNAc-PG are absent in the PA14_56030::Tn mutant. Shown are the extracted ion chromatograms (EICs) of NPLC/MS in the negative ion mode.

To search for the GlcNAc-PG deacetylase, we queried the PA14 genomic database and found six genes annotated as deacetylases. Among them, four have been well characterized with identified substrates, including LpxC (35, 36). However, the other two putative deacetylases, with locus ID PA14_44830 and PA14_56040 (*gpgD*) (Table 2), have not been experimentally characterized and their substrates have remained unknown.

**Table 2.**
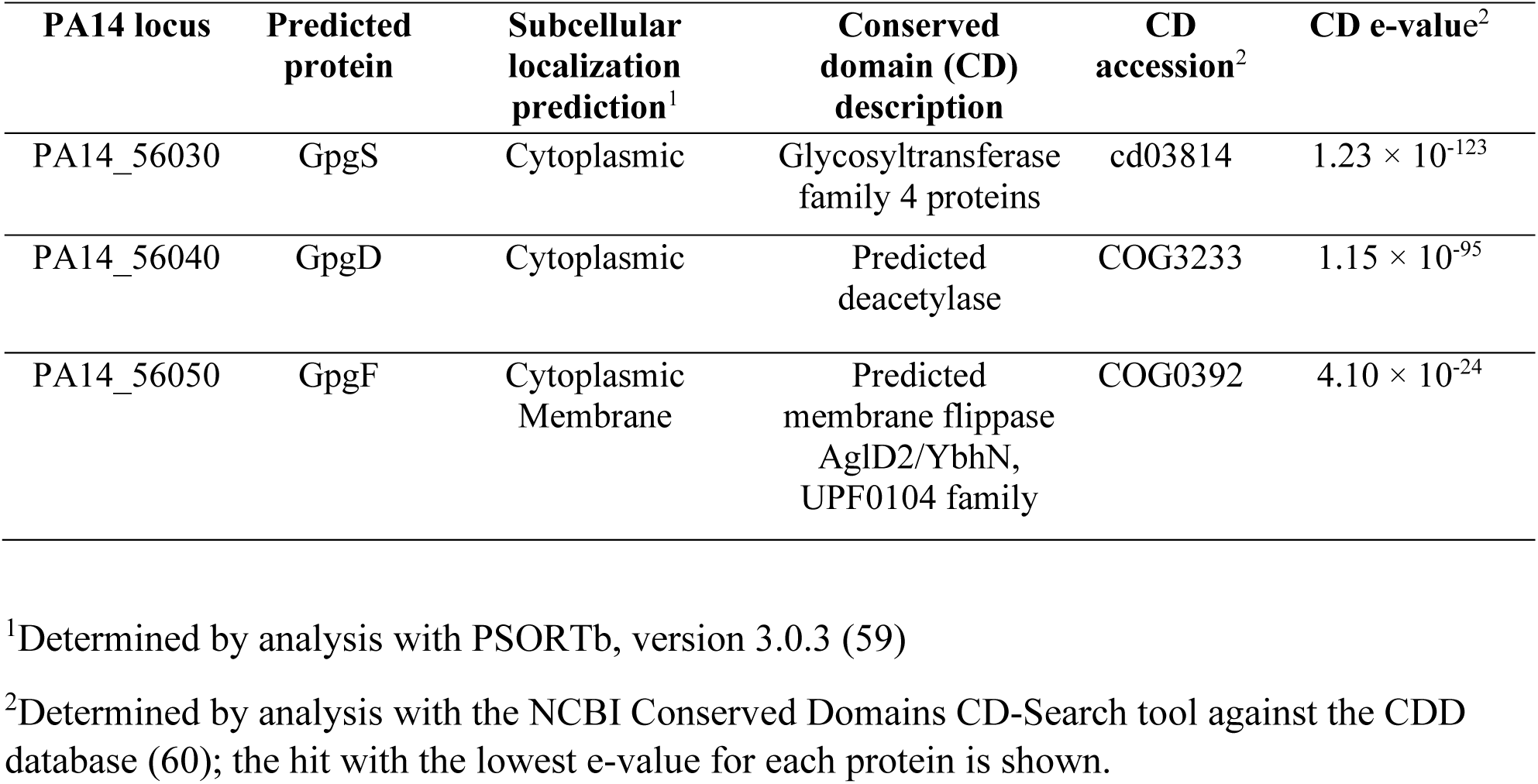
Bioinformatic Analysis of Function and Localization of GpgS, GpgD and GpgF proteins.

To determine whether GlcNAc-PG is a substrate of PA14_44830 and/or PA14_56040, we performed lipidomic analysis of their respective transposon mutants from a previously reported mutant collection (22). As shown in Fig. 2B, PA14_44830 mutant contains both GlcN-PG and GlcNAc-PG, with their relative levels similar to the wild-type (WT), suggesting that PA14_44830 is not a GlcNAc-PG deacetylase. By sharp contrast, the PA14_56040 (*gpgD*) mutant lacks GlcN-PG, but contains drastically elevated GlcNAc-PG (Fig. 2C), strongly supporting that GlcNAc-PG and GlcN-PG are the substrate and product of GpgD, respectively. The identification of the accumulated GlcNAc-PG (Fig. 3A) is further confirmed by MS/MS (Fig. 3B). The complete depletion of GlcN-PG in the PA14_56040 (*gpgD*) mutant also indicates that GpgD is the sole deacetylase responsible for the conversion of GlcNAc-PG to GlcN-PG.

**Figure 3.**
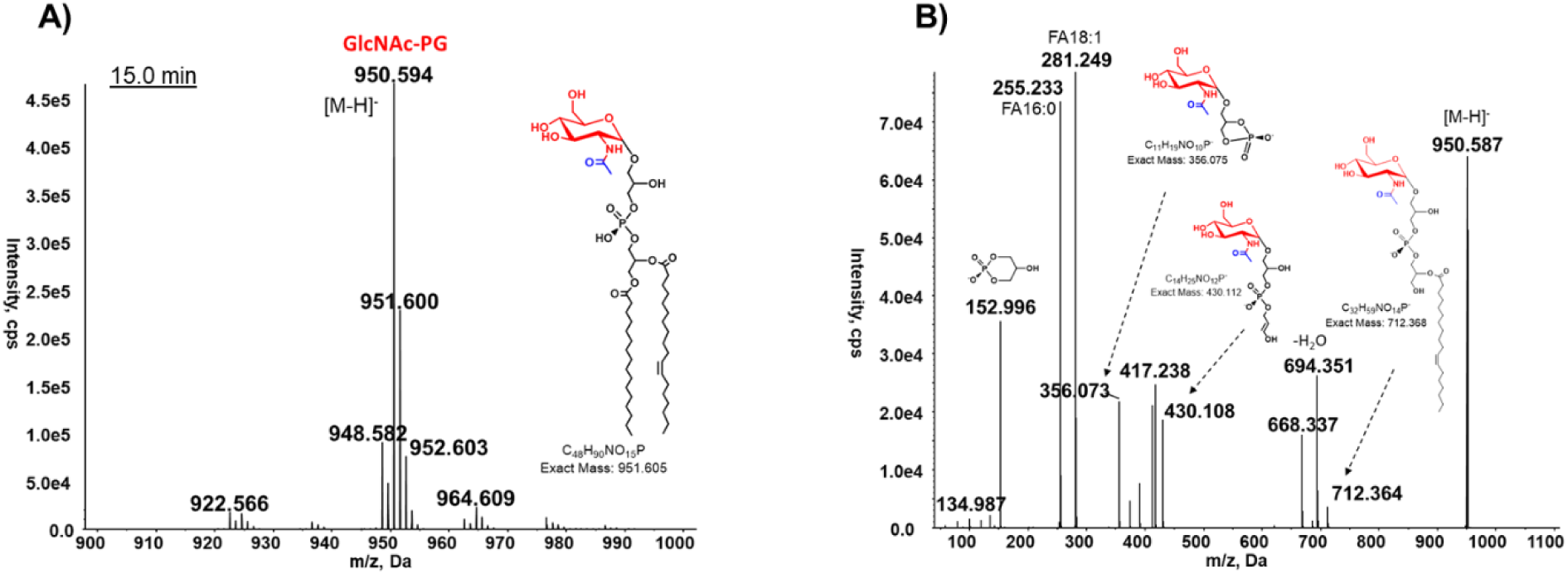
MS analysis of GlcNAc-PG accumulated in the PA14_56040 (*gpgD*)::Tn mutant. A) Negative ion mass spectrum of [M-H]^-^ ion at *m/z* 950.6 for GlcNAc-PG (16:0/18:1). B) MS/MS of [M-H]^-^ ion at *m/z* 950.6 produces fragment ions consistent with GlcNAc-PG (16:0/18:1)

PA14_56030 (*gpgS*) is encoded upstream of *gpgD* and is predicted to be a glycosyltransferase (Table 2). Its predicted function and genomic position strongly suggest that GpgS is likely the glycotransferase responsible for transferring the GlcNAc group onto PG to produce GlcNAc-PG. Indeed our lipidomic analysis shows that the PA14_56030 (*gpgS)*::Mar2xT7 mutant lacks both GlcNAc-PG and GlcN-PG (Fig. 2D).

Overall, the lipidomic analysis of the *gpgD* and *gpgS* transposon mutants reveals that GlcN-PG biosynthesis involves two steps: GpgS first transfers the GlcNAc group to PG to form GlcNAc-PG, which is then deacetylated by GpgD to produce GlcN-PG (Fig. 4). Although not experimentally verified, UDP-GlcNAc is likely the GlcNAc donor based on its utilization in other GlcNAc-modifications (39, 40).

**Figure 4.**
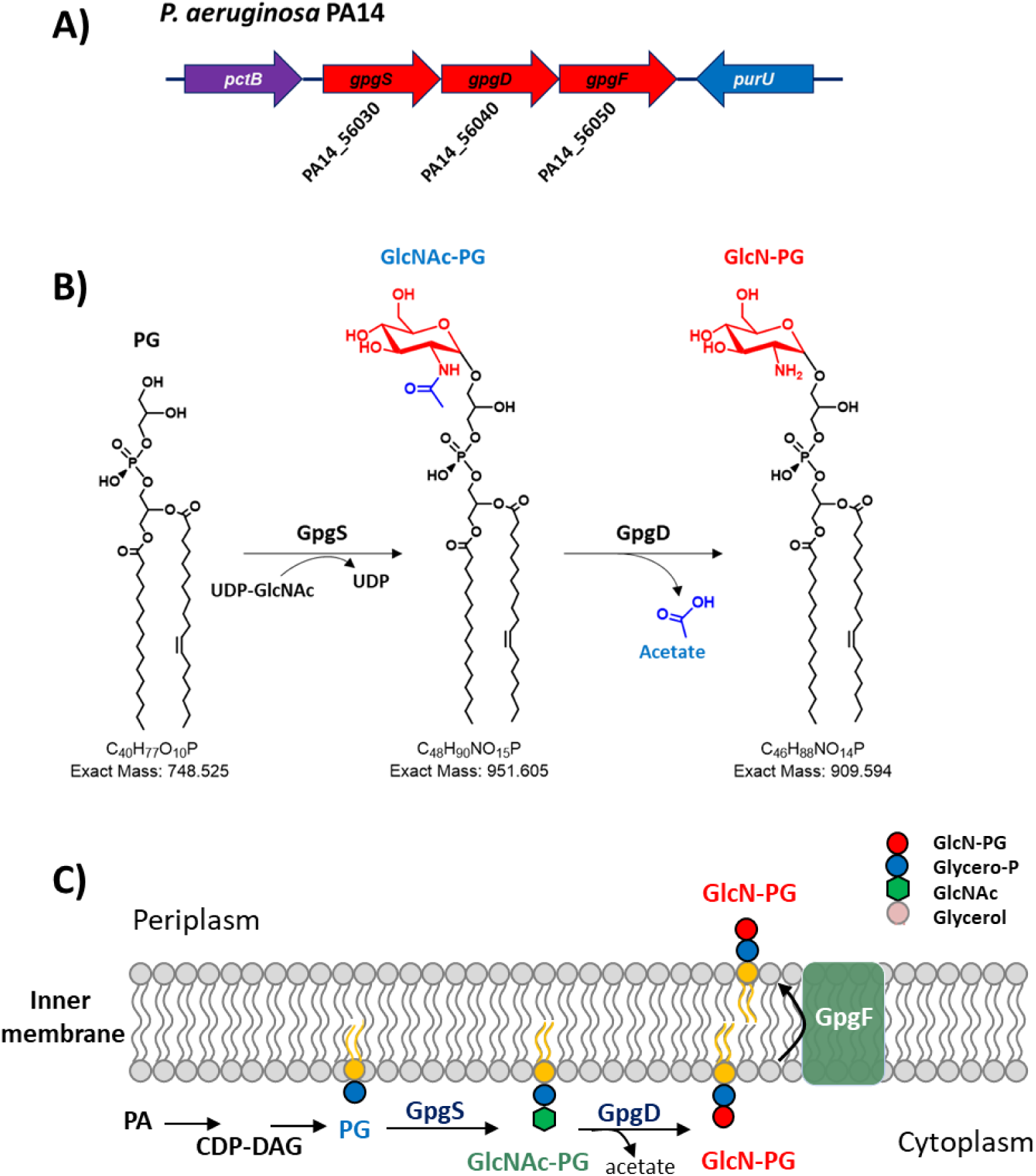
Proposed biosynthesis and translocation of GlcN-PG in *P. aeruginosa*. A) Diagram of the *gpgSDF* predicted operon and its neighboring genes in *P. aeruginosa* PA14. B) The biosynthesis of GlcN-PG from PG involves two enzymatic steps. First, PG is modified by GlcNAc by a glycosyltransferase (PA14_56030, GpgS), presumably using UDP-GlcNAc as the sugar donor. Second, GlcNAc-PG is de-acetylated by a deacetylase (PA14_56040, GpsD) to produce GlcN-PG. C) Cartoon illustration of the proposed biosynthesis and membrane translocation of GlcN-PG in *P. aeruginosa*. The localization of GpgSDF has not been experimentally determined.

We performed complementation of PA14 *gpgD*::Tn mutant, for which a pMQ72 derivative with *gpgD* under the control of an arabinose inducible promoter was constructed (Fig. 5A). Lipidomic analysis showed that arabinose-induced complementation of PA14 *gpgD*::Tn resulted in the production of GlcN-PG (Fig. 5B-C). The complementation of PA14 *gpgS*::Tn mutant remains to be demonstrated. We generated pMQ72-Tet-*gpgS* by cloning the *gpgS* gene in a similar vector system to *gpgD* but have been unable to obtain colonies of the PA14 *gpgS*::Tn mutant bearing pMQ72-Tet-*gpgS* despite trying various approaches, including electroporation, chemical transformation, and conjugation.

**Figure 5.**
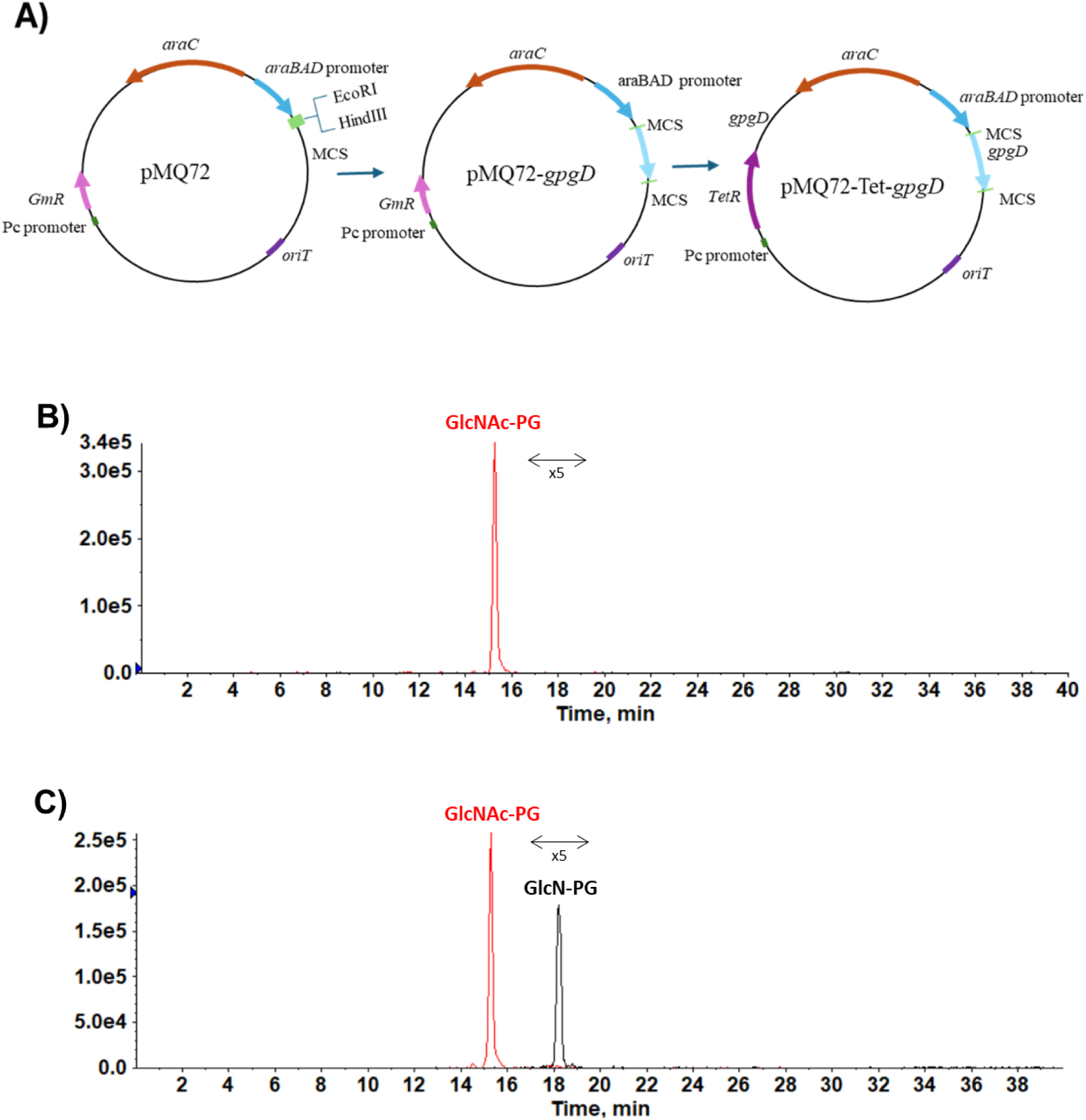
Complementation of the *gpgD*::Tn mutant of PA14 with *gpgD in trans* restores GlcN-PG production. A) Construction of a pMQ72 derivative with *gpgD* under the control of an arabinose-inducible promoter and containing a tetracycline resistane cassette. B) GlcN-PG is absent without arabinose induction. C) GlcN-PG is produced with arabinose (1%) induction. For better illustration, the GlcN-PG peak is magnified by 5-fold.

PA14_56050 (renamed *gpgF*), predicted to be a flippase (Table 2), is downstream of *gpgS* and *gpgD*. To experimentally demonstrate its flippase function will require the development of an assay that can monitor the membrane translocation of GlcN-PG (41, 42). Preliminary lipidomic analysis of the PA14_56050:Tn mutant shows that it contains both GlcN-PG and GlcNAc-PG (Fig. S1). Compared to WT, the level of GlcN-PG seems significantly lower (by several folds) in the PA14_56050::Tn mutant, whereas GlcNAc-PG is slightly increased. The mechanism of the effects of GpgF on GlcN-PG and GlcNAc-PG production remains to be investigated. The presence of GlcN-PG in the *gpgF* mutant seems to suggest that GpgD functions cytosolically (Fig. 4), a hypothesis assuming that GpgF is the only flippase for GlcNAc-PG or GlcN-PG. The function of *gpgSDF* in PA14 is analogous to the function of *hexSDF* in *C. difficile*, which is involved in the synthesis of a novel glycolipid, aminohexosyl-hexosyldiradylglycerol (HNHDRG) (34, 43). It is worth noting that the localization of HexSDF was not experimentally determined; both intracellular and extracellular activities of HexD were proposed (34).

### *P. aeruginosa gpgSDF* is sufficient for the synthesis of GlcNAc-PG and GlcN-PG in *E. coli*

*E. coli* contains PG as one of its most abundant phospholipids but does not have *gpgSDF* and does not produce GlcNAc-PG or GlcN-PG (28, 31, 44). To confirm the biochemical activities of *gpgSDF*, the plasmid pWO-GEM-T with the entire *gpgSDF* predicted operon of PA14 was constructed from pGEM-T vector and was then transformed into *E. coli* DH5α (Fig. 6A). Lipidomic analysis detected both GlcNAc-PG and GlcN-PG (Fig. 5B-D) in *E. coli* with pWO-GEM-T, confirming that *gpgSDF* possesses the expected glycotransferase and deacetylase activities. The identification of GlcNAc-PG and GlcN-PG produced in *E. coli* are supported by exact measurement and MS/MS. It is worth noting that the acyl chain profiles of the GlcNAc-PG and GlcN-PG molecular species heterologously produced in *E. coli* are slightly shorter (about 2 carbon atoms less) than those in *P. aeruginosa.* As expected, the expression of *gpgS* alone in *E. coli* resulted in the production of GlcNAc-PG, but not GlcN-PG (Fig. S2).

**Figure 6.**
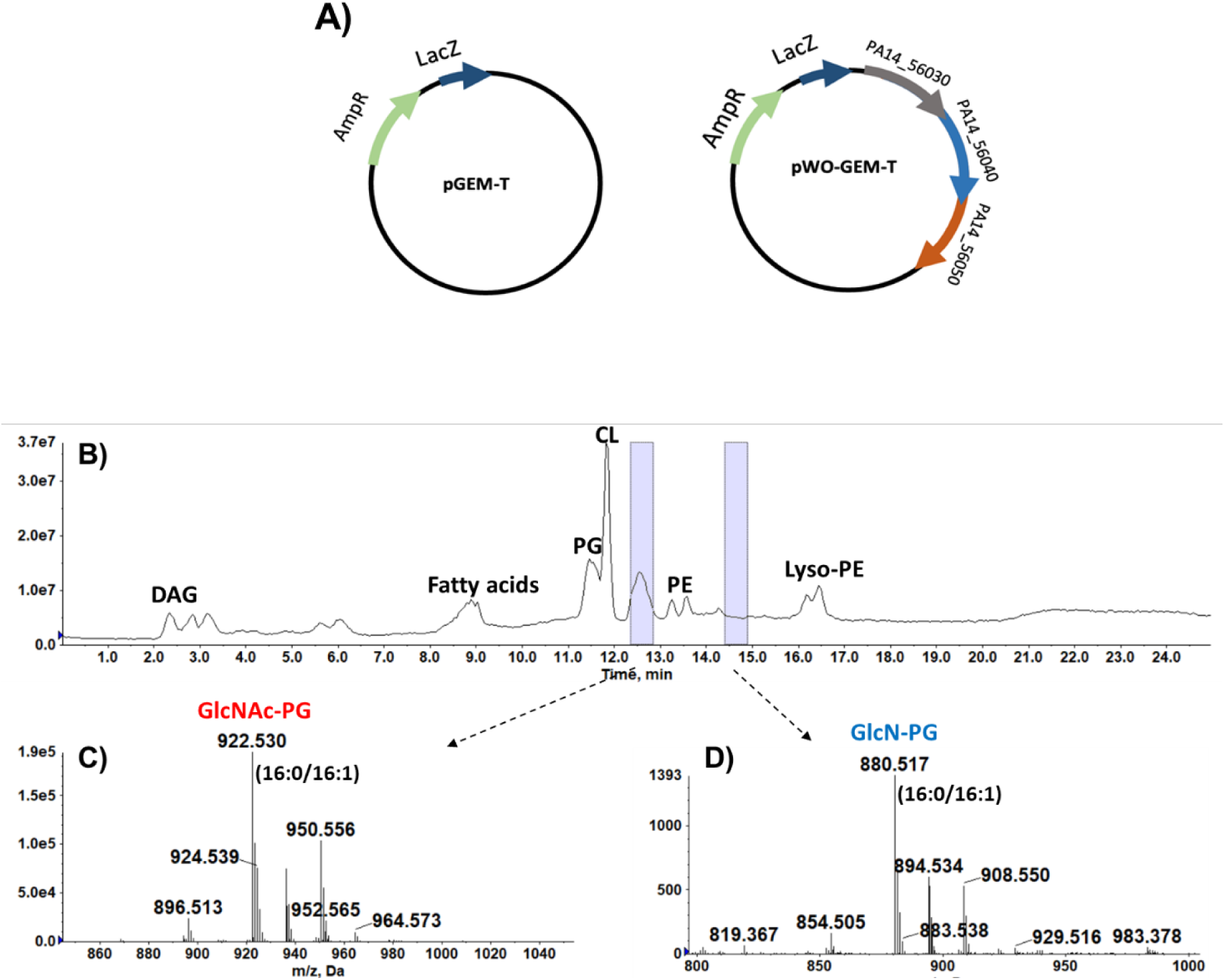
The heterologous expression of the PA14 *gpgSDF* genes produces GlcNAc-PG and GlcN-PG in *E. coli*. A) The plasmid pWO-GEM-T with the *gpgSDF* genes of PA14 was constructed from pGEM-T vector and was then transformed into *E. coli* DH5α. B) Total ion chromatogram of the normal phase LC/MS in the negative ion mode of the lipid extract of *E. coli* expressing the pWO-GMT-T. C) Negative ion mass spectrum of the [M-H]^-^ ions of GlcNAc-PG species (appearing at ∼12.5-13.0 min). D) Negative ion ass spectrum of the [M-H]^-^ ions of GlcN-PG species (appearing at ∼ 14.5-15.0 min). The identification of GlcNAc-PG (16:0/16:1) and GlcN-PG (16:0/16:1) is confirmed by MS/MS.

GlcN-PG is much less abundant than GlcNAc-PG in the *gpgSDF-*transformed *E. coli* (Fig. 6B-D), compared to *P. aeruginosa*. It is unknown whether this is due to altered codon usage in *P. aeruginosa* relative to *E. coli*, a non-optimal Shine-Dalgarno sequence for *gpgD* in the heterologous host, or other factors. Further optimization of the translation and activity of GpgD will be needed to improve the conversion of GlcNAc-PG to GlcN-PG to facilitate further studies of these lipids in *E. coli*.

### Identification of GlcN-PG in other *P. aeruginosa* strains

Our bioinformatic analysis indicates that the *gpgSDF* operon is conserved in all sequenced strains of *P. aeruginosa.* Indeed, our lipidomic analysis confirmed the presence of GlcN-PG in three other strains of *P. aeruginosa* (Fig. 7 and Table 3).

**Figure 7.**
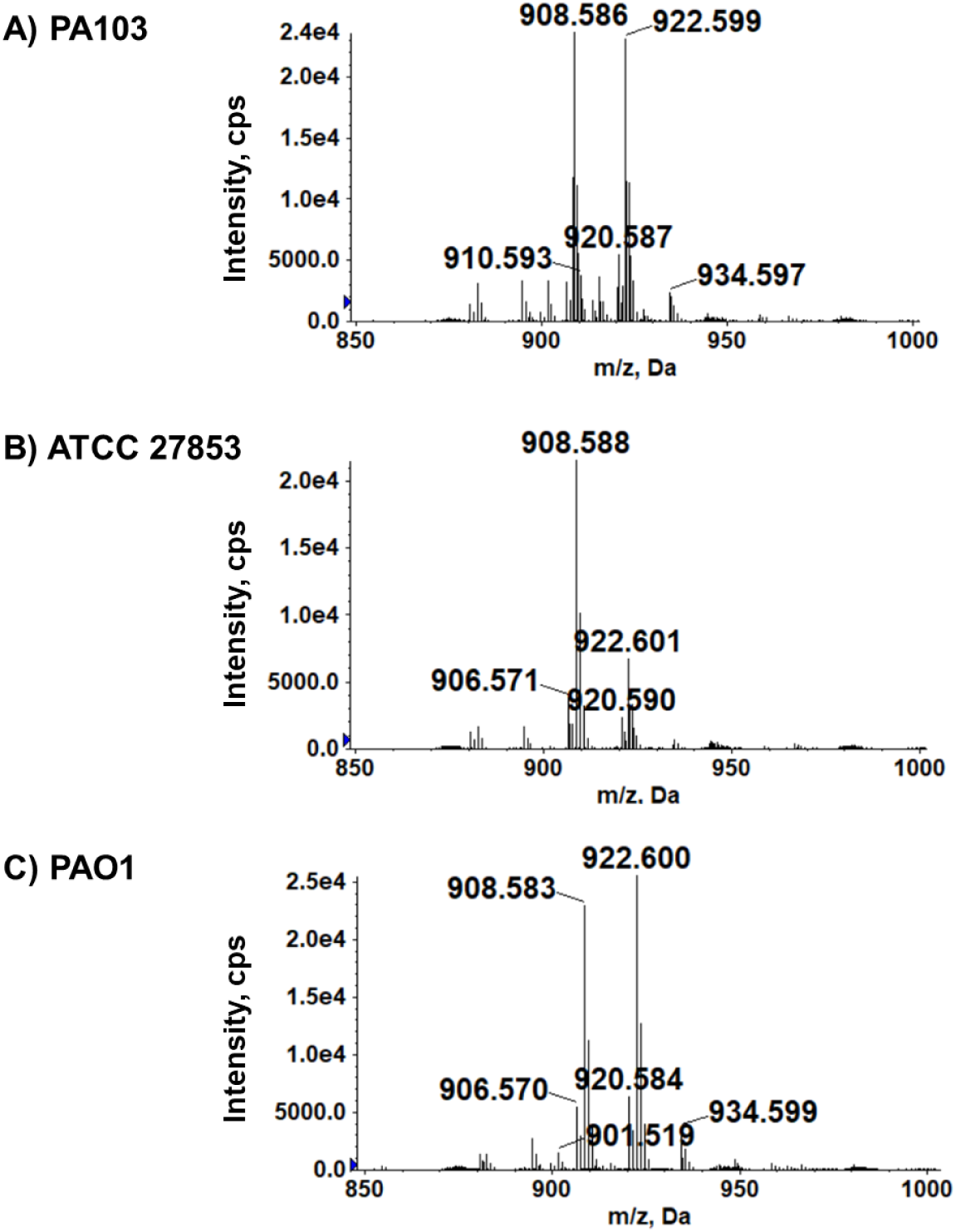
Detection of GlcN-PG in various *P. aeruginosa* laboratory and clinical strains by LC/MS. Shown are the negative ion mass spectra of the [M-H]^-^ ions of GlcN-PG species in A) PA103, B) ATCC 27853, C) PAO1 (Table 2). The identification of GlcN-PG molecular species is confirmed by exact mass measurement and MS/MS.

**Table 3.** Genomic and lipidomic analysis of GlcN-PG in *P. aeruginosa* stains.

| Strains | *gpgSDF* operon (Accession numbers shown for each) |  |  | GlcN-PG |
| --- | --- | --- | --- | --- |
|  | *gpgS* | *gpgD* | *gpgF* |  |
| PAO1 | PA4311<br>(NP_253001.1) | PA4312<br>(NP_253002.1) | PA4313<br>(NP_253003.1) | ✓ |
| PA14 | PA14_56030<br>(YP_792656.1) | PA14_56040<br>(YP_792657.1) | PA14_56050<br>(YP_792658.1) | ✓ |
| PA103 | PA103_5903<br>(WP_003110035.1) | PA103_5904<br>(WP_003112786.1) | PA103_5905<br>(WP_019485776.1) | ✓ |
| ATCC 27853 | RS24835<br>(WP_003101181.1) | RS24840<br>(WP_003101179.1) | RS24845<br>(WP_016852096.1) | ✓ |

## Discussion

Using a combination of lipidomic and bioinformatic approaches, we identified two biosynthetic enzymes that are involved in GlcN-PG synthesis in *P. aeruginosa*. GpgS is a novel glycotransferase that catalyzes the modification of phosphatidylglycerol (PG) with *N*-acetylglucosamine (GlcNAc) to produce GlcNAc-PG, and GpgD is a novel deacetylase that removes the acetyl group of GlcNAc-PG to produce GlcN-PG. The functions of these enzymes were further confirmed by the synthesis of GlcNAc-PG and GlcN-PG from the heterologous expression of *gpgSDF* in *E. coli*.

The membrane distribution of GlcN-PG remains to be determined. *gpgF* is predicted to encode a flippase. Bioinformatic analysis predicts that GpgF has transmembrane spanning helices with inner membrane localization, whereas both GpgS and GpgD lack transmembrane spanning helices and are predicted to be cytoplasmic (Table 2). Thus, the biochemical activities mediated by *gpgSDF* are analogous to MprF which catalyzes Ala-PG synthesis on the cytosolic side of the inner membrane and subsequent translocation of Ala-PG to the periplasmic side of inner membrane (45, 46). In *P. aeruginosa*, it is unknown whether GlcN-PG, if flipped across the inner membrane, would be further transported to the outer membrane via proteins such as the AsmA-like family (47).

The identification of the GlcN-PG biosynthetic genes will enable the study of the biological and pathological functions of GlcN-PG in *P. aeruginosa*. Previously it was observed that GlcN-PG was dramatically increased under acidic conditions (32) for *Bacillus megaterium* and at stationary phase (17) for *P. aeruginosa*, suggesting the production of GlcN-PG is associated with coping with environmental and nutritional stresses. Previous studies have shown the *gpgS* gene (PA4311 in PAO1 strain) to be involved with phenazine production (48), differentially regulated under oxidative stress (49), implicated in exopolysaccharide alginate production (50), and up-regulated in acute and chronic infections (51).

The replacement of anionic PG with zwitterionic GlcN-PG is expected to lower the net negative charge of the bacterial membrane and thus may help confer resistance to cationic antimicrobials. In gram-positive *C. difficile*, GlcN-modification of a glycolipid impacts resistance to cationic antibiotics including daptomycin and bacitracin (34). It is plausible that GlcN-PG plays a similar role in defending against cationic compounds that are toxic to *P. aeruginosa*.

Remarkably, the lipidome of *P. aeruginosa* is more diverse than *E. coli* which has been the model organism for studying lipid composition and metabolism in Gram-negative bacteria. In particular, *P. aeruginosa* contains a multitude of zwitterionic phospholipids, including GlcN-PG, while *E. coli* has only one zwitterionic phospholipid (PE) in addition to two major anionic phospholipids (PG and CL) (28, 31, 44) (Fig. 8). The lipid diversity of *P. aeruginosa* reflects its relatively large genome size (52, 53) and likely contributes to its unusual metabolic flexibility and membrane adaptability. The elucidation of the functions of GlcN-PG and other *P. aeruginosa* zwitterionic phospholipids will be important for fully understanding the molecular mechanisms underlying the unique membrane properties, environmental resilience and adaptability, and high propensity for antibiotic resistance development of *P. aeruginosa*.

**Figure 8.**
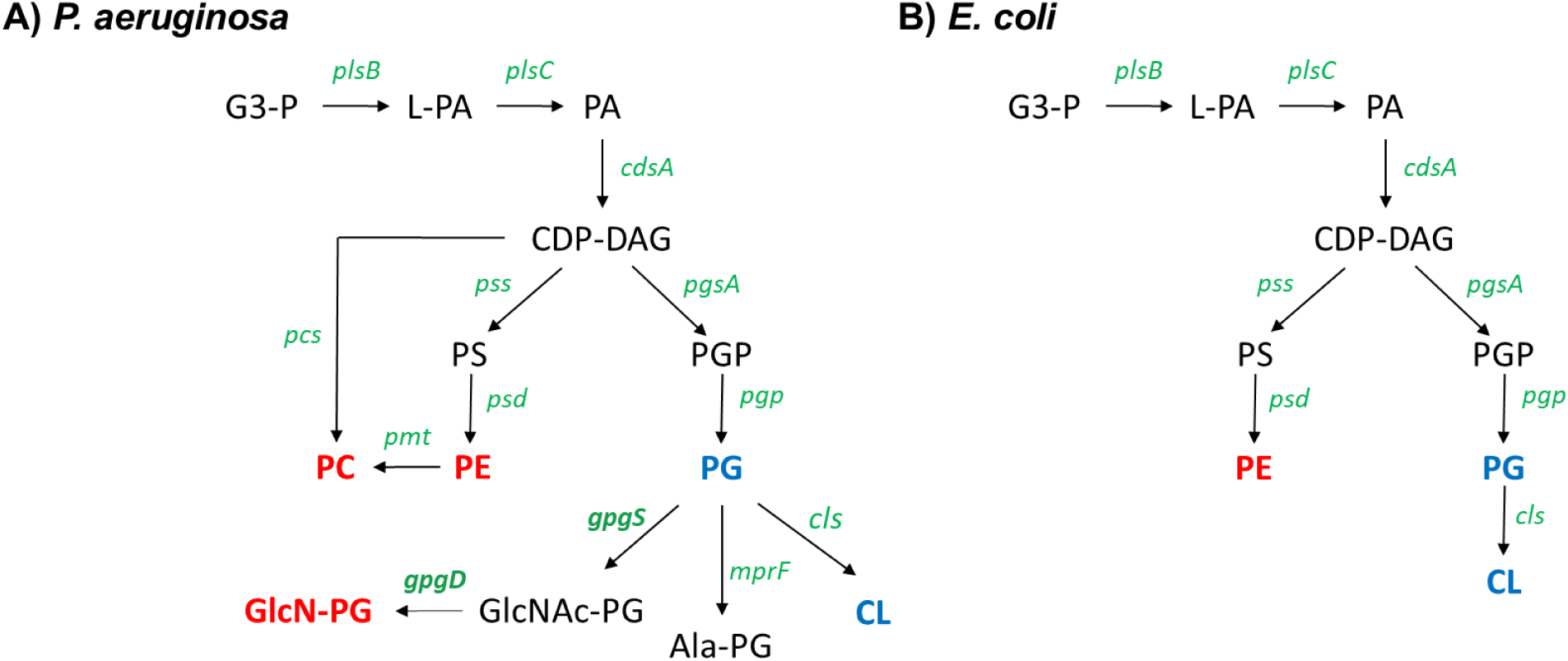
Comparison of phospholipid biosynthetic pathways in *P. aeruginosa* and *E. coli*. *P. aeruginosa* possesses more complex phospholipids than *E coli*. Major zwitterionic phospholipids (red) and anionic phospholipids (blue) are in bold fonts. Their biosynthetic genes are green. The synthesis of GlcN-PG from PG in *P. aeruginosa* (the subject of this study) consists of two enzymatic steps catalyzed by GpgS and GpgD. Abbreviations: G3-P, glycerol-3-phosphate; L-PA, lyso-phosphatidic acid; PA, phosphatidic acid; CDP-DAG, CDP-diacylglycerol; PS, phosphatidylserine; PE, phosphatidylethanolamine; PC, phosphatidylcholine; PGP, phosphatidylglycerol-3-P; PG, phosphatidylglycerol; Ala-PG, alanyl-phosphatidylglycerol; CL, cardiolipin.

## Acknowledgements

This work was supported by grants R01AI148366 and R01AI178692 from the National Institutes of Health to K.L.P. and Z.G. and the Cecil H. and Ida Green Chair in Systems Biology Science to K.P. The authors would like to thank the Dillon lab at the University of Texas at Dallas for providing the PA103 and ATCC 27853 strains of *P. aeruginosa*, Dr. Peter Jorth for providing pMQ72, and Dr. Trusha Parekh for providing *E. coli* S17.

## Supplemental Information

**Figure S1.**
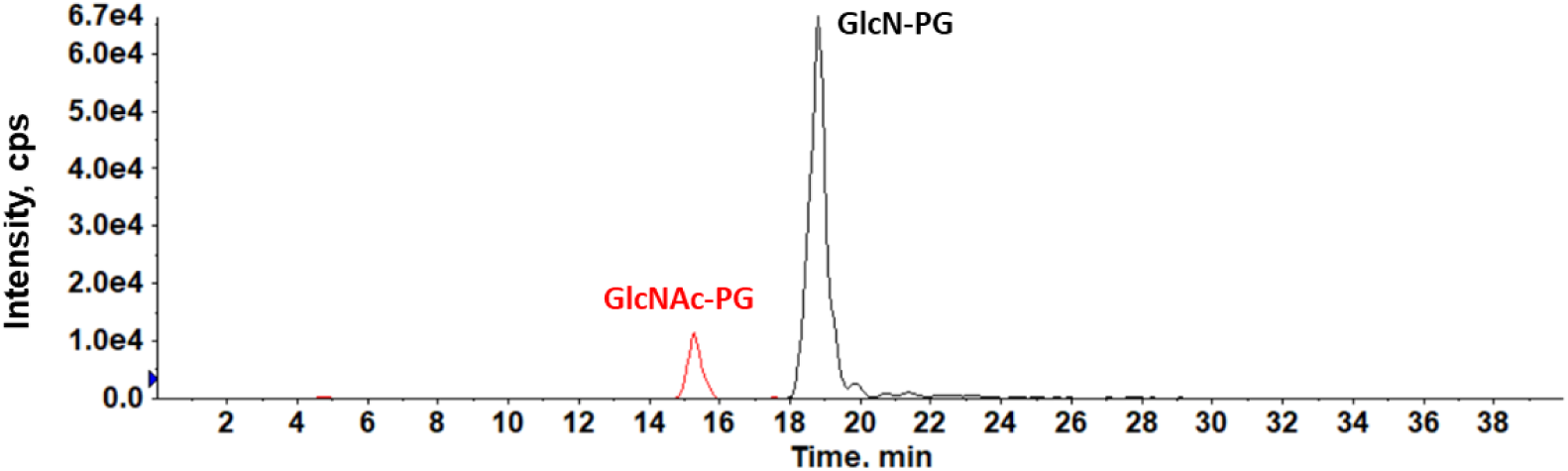
LC/MS analysis of GlcNAc-PG and GlcN-PG in the PA14_56050 (*gpgF*) mutant of *P. aeruginosa*. PA14_56050 (*gpgF*) is bioinformatically predicted to be a lipid flippase. Lipidomic analysis of the PA14 56050::Tn mutant shows that it contains both GlcN-PG and GlcNAc-PG. Compared to WT, the level of GlcN-PG is significantly lower (roughly by several folds), whereas GlcNAc-PG is slightly increased in the PA14 56050::Tn mutant. The presence of GlcN-PG (at a much higher level than GlcNAc-PG) in the *gpgF* mutant suggests that GpgD functions cytosolically, assuming that GpgF is the only flippase for GlcNAc-PG or GlcN-PG.

**Figure S2.**
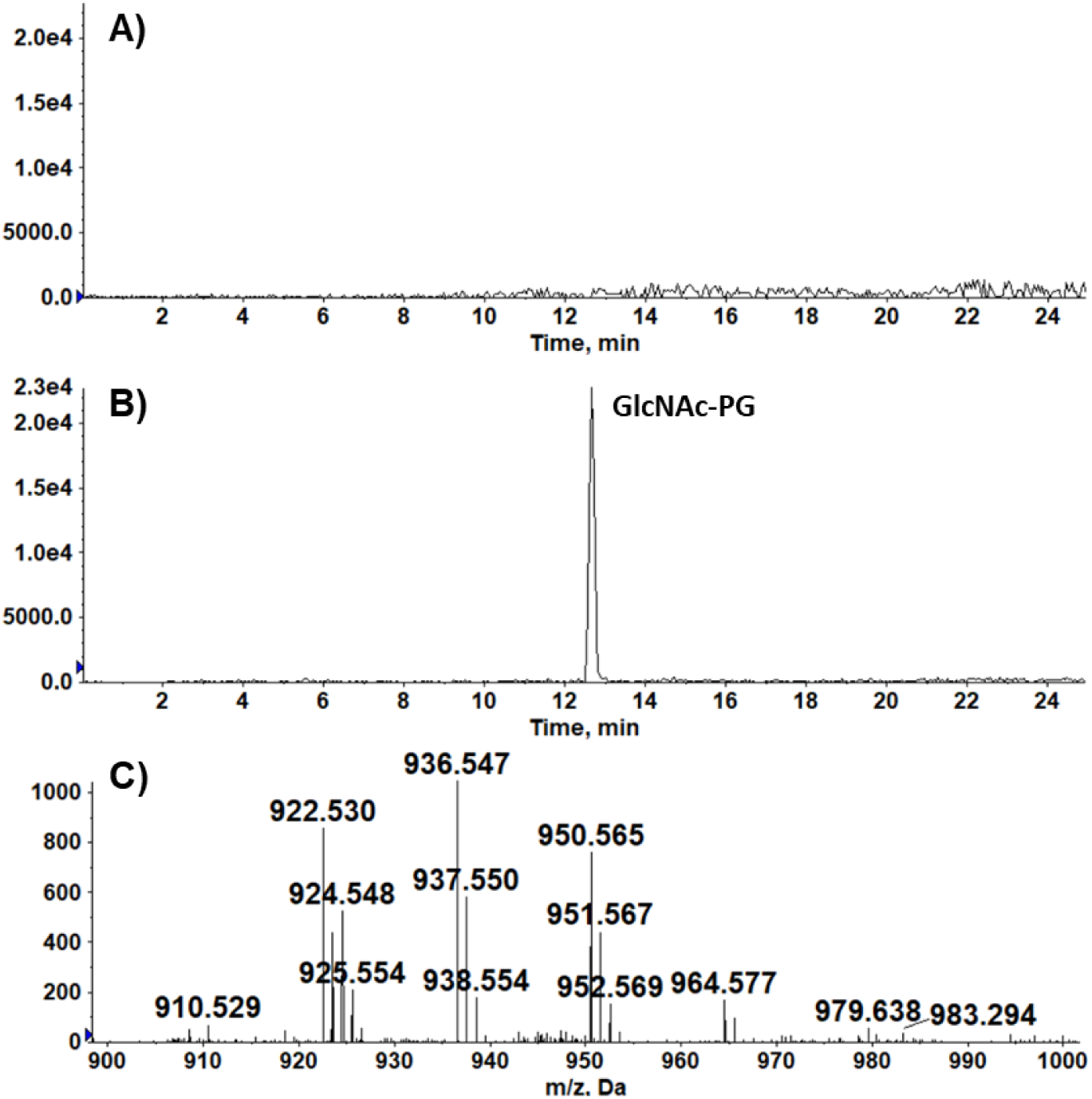
Heterologous synthesis of GlcNAc-PG in *E. coli* by expressing PA14_56030 (*gpgS*) from *P. aeruginosa*. A) *E. coli* DH5a lacks both GlcN-PG and GlcNAc-PG. B) GlcNAc-PG, but not GlcN-PG, is produced by expressing PA14_56050 (GpgS) in *E. coli* DH5a. C) Negative ion mass spectrum of [M-H]^-^ ions of GlcNAc-PG species produced by expressing PA14_56050 (GpgS) in *E. coli* DH5a.

**Table S1.**
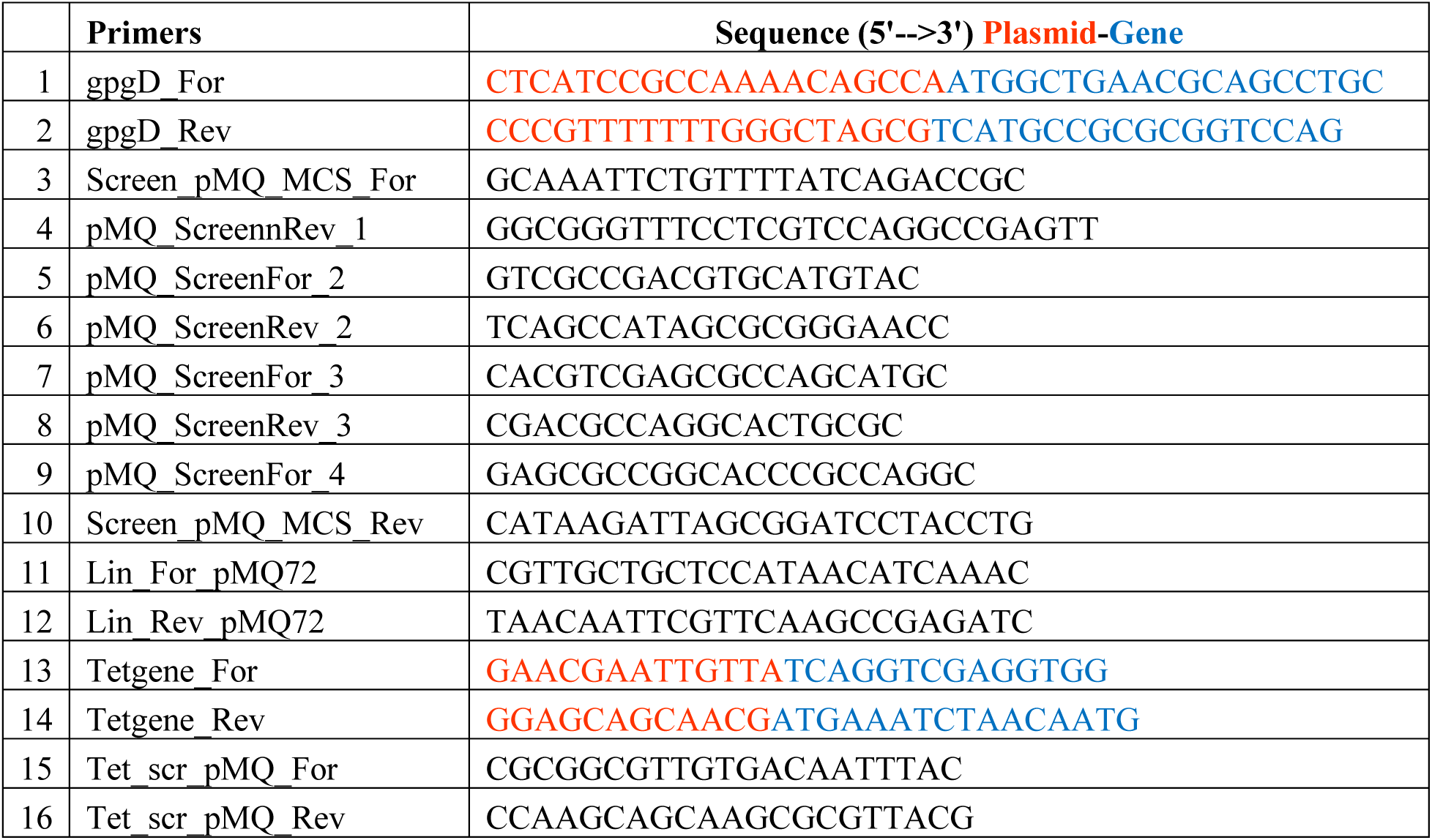
Primers for *gpgD* complementation in PA14 *gpgD*::Tn mutant.

